# The long and the short of it: Nanopore based eDNA metabarcoding of marine vertebrates works; sensitivity and specificity depend on amplicon lengths

**DOI:** 10.1101/2021.11.26.470087

**Authors:** Karlijn Doorenspleet, Lara Jansen, Saskia Oosterbroek, Pauline Kamermans, Oscar Bos, Erik Wurz, Albertinka Murk, Reindert Nijland

## Abstract

To monitor the effect of nature restoration projects in North Sea ecosystems, accurate and intensive biodiversity assessments are vital. DNA based techniques and especially environmental DNA (eDNA) metabarcoding is becoming a powerful monitoring tool. However, current approaches are based on genetic target regions <500 nucleotides, which offer limited taxonomic resolution. We developed a method for eDNA metabarcoding, based on nanopore sequencing of a longer amplicon, enabling improved identification of fish species. We designed a universal primer pair targeting a 2kb region of fish mitochondria, and compared it to the commonly used MiFish primer pair that targets only ∼170bp. *In sillico* and mock community testing showed that the 2kb fragments improved the accurate identification of genetically closely related species. eDNA was amplified, and sequenced using the Oxford Nanopore MinION in combination with the sequence read processing pipeline Decona. Analyzing eDNA from a North Sea aquarium showed that sequences from both primer pairs could be assigned to most species, but both approaches also identified unique species in the aquarium eDNA. Next, both primer pairs were used on multiple eDNA samples from the North Sea. Here, similar location specific fish communities were obtained from both approaches. More species were identified through the MiFish approach in the field samples. Interestingly, this difference was not observed in the aquarium, suggesting that 2kb fragment based metabarcoding potentially detects more recent occurrences of animals. This new method has the potential to improve and expand the molecular toolbox for eDNA based monitoring approaches.

## Introduction

North Sea fish populations are sensitive to disturbances such as fisheries, nutrient run-off and increasing sea water temperatures (Andersen et al., 2020; Capuzzo et al., 2018; Hofstede et al., 2010; Krehenwinkel et al., 2019; O’Brien et al., 2019). Combined management strategies such as reduced fishing (Couce et al., 2020), designation of marine protected areas (MPA), and placing artificial hard substrates such as off-shore wind parks are suggested to facilitate rehabilitation of the North Sea ecosystem (Claudet, 2018; Degraer et al., 2020; Didderen et al., 2019; Kamermans et al., 2018). To understand how North Sea fish population dynamics are affected by these strategies, development and validation of methods that map fish population diversity and density is crucial. Conventional marine fish biomonitoring practices largely rely on destructive methods that involve netting and trapping (Daan et al., 2005; Reiss et al., 2010). These methods are costly, time-consuming and require expert taxonomic visual identification skills (Mateos-Rivera et al., 2020; Teletchea, 2009). In addition, conventional methods have limited sampling efficiencies and may be disruptive to the environment (Eggleton et al., 2018). Thus, it is crucial to develop precise and non-invasive biomonitoring solutions that are also time and cost efficient (Goodwin et al., 2017).

Environmental DNA (eDNA) based fish species identification has gained substantial attention in the last decade, as it can detect the presence of fish species based on a small amount of DNA present in seawater. It has been shown to be highly sensitive for non-indigenous species detection (Ficetola et al., 2008) and identification of spawning and migration patterns (Thalinger et al., 2019). Short amplicon eDNA metabarcoding has become an increasingly popular tool to perform fish community assessment for identification of ecological relevant fish species from an array of ecosystems (Deiner et al., 2017; Miya et al., 2015a; Ruppert et al., 2019a; Taberlet et al., 2012; Thomsen et al., 2012). Also in the North Sea, eDNA metabarcoding results from a sampling effort close to fykes showed to be comparable to the fyke catches themselves. (Bleijswijk et al., 2020).

The standardization of eDNA metabarcoding as marine monitoring strategy is still under development. Species-specific differences occur in e.g. degree of skin cell shedding, degradation rates vary depending on temperature and season, and unknown dilution factors depending on currents all make quantification of the results challenging (Beng & Corlett, 2020; Lacoursière-Roussel et al., 2016; Sassoubre et al., 2016; Seymour et al., 2018). The sample preparation, metabarcoding technique and workflow will determine the quality of the results and thus the species detection quality and possible biases (Beng & Corlett, 2020; van der Loos & Nijland, 2020). Important steps in the protocol include decisions about methods of sampling and DNA extraction (Bessey et al., 2020; Hunter et al., 2019), primer and PCR settings (Doi et al., 2019; Sard et al., 2019; Zhang et al., 2020), sequencing technology (Egeter et al., 2020; Singer et al., 2019; Truelove et al., 2019), post-sequencing data handling (Santos et al., 2020a) and reference databases used (Hestetun et al., 2020; McGee et al., 2019).

Especially choice of primer pair and targeted DNA region are crucial for successful fish detection with eDNA (Beng & Corlett, 2020). Several universal fish primers are described that mostly target regions of the mitochondrial genome as there is a high copy number of this genome per cell (Schon, 2000). The most used primers target different short regions from 100 to 500 nucleotides of the 12S rRNA (Miya et al., 2015a; Riaz et al., 2011; Taberlet et al., 2018) 16S rRNA (DiBattista et al., 2017; Evans et al., 2016), cytochrome B (Thomsen et al., 2012) and COI gene (Balasingham et al., 2018). Although primers targeting short 12S regions are the most commonly used and considered as a standard (Shu et al., 2020), longer target amplicon size yield higher taxonomic resolution (Zhang et al., 2020). Also the use of multiple primer pairs are suggested to increase taxonomic resolution (Evans et al., 2016; Miya et al., 2015a; Zhang et al., 2020) and has been shown to increase species level detectability in lakes (Sard et al., 2019). Thus, using longer fragments and multiple marker sets will enhance the species resolution and specificity.

Long read sequence analysis with commonly used Illumina platforms is not possible due to its abilities to sequence with high accuracy but with a read length maximum of 500bp (Tan et al., 2019). Fortunately, third generation sequencing as available from Oxford Nanopore Technologies (ONT) and Pacific Biosciences enables the generation of ultra-long sequences (Bleidorn, 2016). This can be used for eDNA studies that are based on primer pairs targeting longer regions, covering several mitochondrial marker genes. This long amplicon approach was demonstrated to be successful in microbial metabarcoding studies, and increased taxonomic resolution to species level and beyond (Johnson et al., 2019; Shin et al., 2016). Historically, the main limitation of nanopore sequencing was the relatively large error rate of 5 to 10% (Jain et al., 2015). This error rate can be overcome with bioinformatics tools to generate reliable consensus sequences and thus increase sequence accuracy (Baloğlu et al., 2020; Carradec et al., 2020; Egeter et al., 2020). To our knowledge, a bioinformatics pipeline was not yet available to generate such consensus sequences from raw sequence data in multiplexed metabarcoding experiments. Such a pipeline would greatly facilitate the development of long read metabarcoding in marine molecular biomonitoring.

This study investigates the possibilities of long read metabarcoding of eDNA with a newly designed fish and other vertebrates specific primer pair targeting a 2kb target fragment containing parts of both the 12S and 16S mitochondrial rRNA genes, using Oxford Nanopore MinION sequencing. To increase species identification accuracy, we developed a sequence data processing pipeline. The identification resolution of the primer pair was tested *in silico* on several genetically similar species. The 2kb primer pair was further evaluated on field samples, and compared with the commonly used universal short read MiFish primer pair targeting a ∼170 bp region of the 12S mitochondrial rRNA (Miya et al., 2015b). We compared the sensitivity and specificity of both primer pairs both using a North Sea aquarium with a known species composition and field samples from different locations and habitat types in the North Sea.

## Materials and Methods

### Sample collection

Samples were collected on three separate sampling sites; the North Sea “Ray Reef” aquarium from the Dolfinarium, Harderwijk, The Netherlands; the North Sea on locations in a transect from Gemini Wind Park to Borkum Reef; and in three different shipwrecks (figure 1).

**Figure 1:**
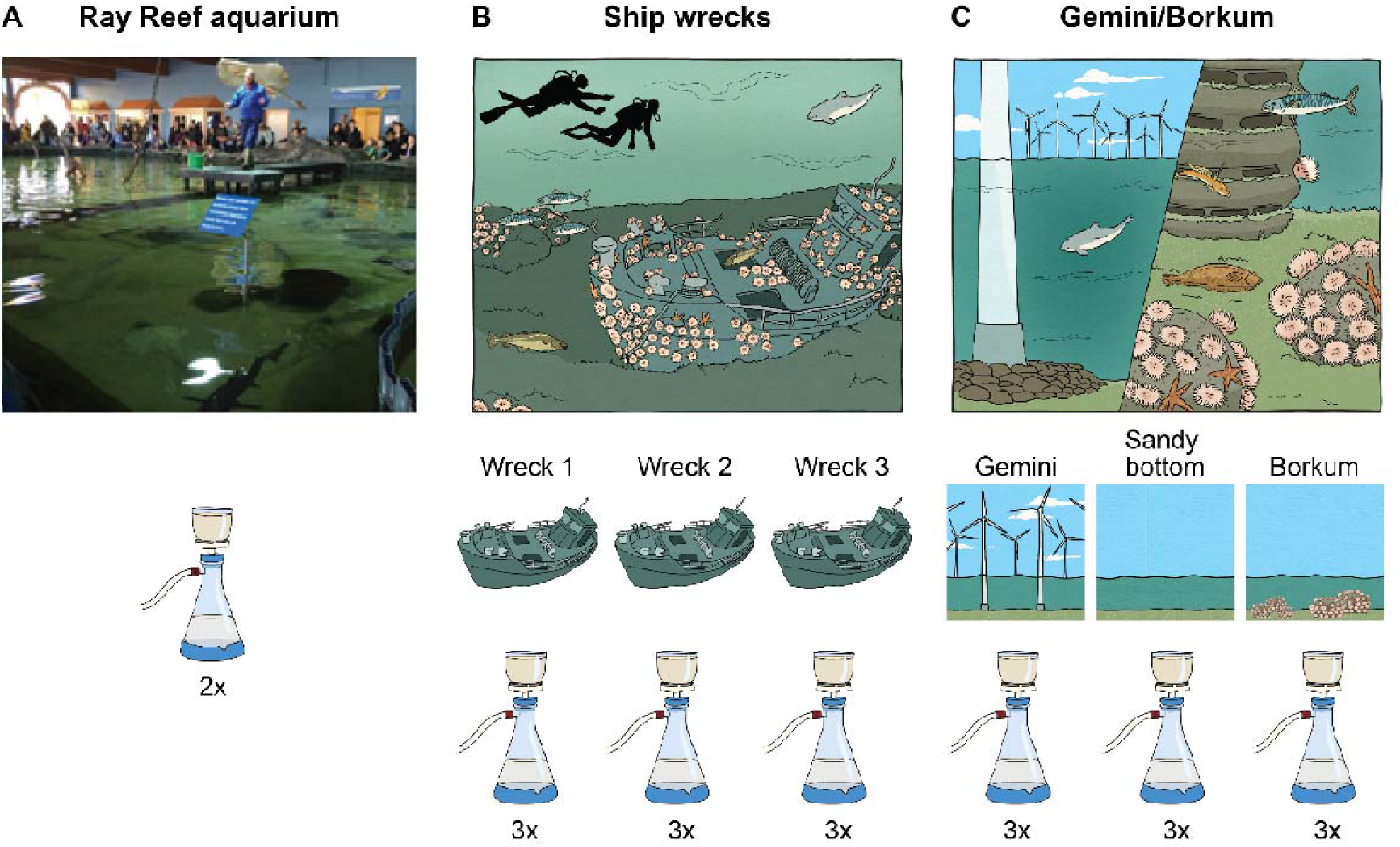
Sampling design of A) North Sea “Ray Reef” aquarium, Dolfinarium, Harderwijk, the Netherlands B) North Sea ship wrecks with three different ship wreck locations and C) Borkum/Gemini where samples were taken in Gemini, halfway between Gemini and Borkum on a sandy bottom and Borkum reef grounds.

For eDNA samples from the ray reef aquarium, two 1L water samples were collected just under the water surface using a 1L plastic container pre-sterilized with bleach. Demineralized tap water was used as negative control. The ray reef aquarium has a volume of 200 m^3^ artificial seawater and represents a North Sea reef that contain predominantly sharks and rays with in total 22 species. The water temperature was 13°C, the salinity at 32.0‰ and the pH at 8.2 at the day of sampling. From Borkum Reef Grounds/Gemini Wind Park, samples were collected from inside Gemini Wind Park (54.0109, 6.0781), halfway between Gemini Wind Park and the Borkum Reef Grounds on sandy substrate (53.8645, 6.2145) (Sandy bottom) and at Borkum Reef Grounds (53.7016, 6.3467). All samples were taken at slack tide during neap tides. Data on environmental parameters at the North Sea sampling locations were retrieved from the Copernicus Marine Service’s Data Portal. Salinity varied in July between 31.8‰ and 34.4‰, temperature 15.7-18.6°C, and pH 7.9-8.1. Three 1L replicates were collected at each location by sampling seawater using 2.5L Niskin bottles at 0.5-1m above the seafloor on 2^nd^ July (figure 1C). Demi water was used as negative control.

For the ship wrecks, samples were collected around three different ship wrecks in the North Sea while SCUBA diving: wreck 1 (55.1821 N, 03.4446 E using the WGS84 reference system), wreck 2 (55.2609, 03.5117) and wreck 3 (55.0774, 02.5087). At each sample location, three replicates of approximately 2L of seawater were taken by pumping (figure 1B, dx.doi.org/10.17504/protocols.io.6yfhftn). Tap water was used as negative control.

All samples were immediately filtered using Thermo Scientific Nalgene Rapid-Flow sterile disposable Filter Units CN (Cellulose nitrate) with a pore size of 0.8µm. Filters were then individually placed in 2mL screwcap Eppendorf tubes. The tubes were prefilled with 400µL Zymo DNA/RNA shield (Zymo, USA) preservative. Samples were immediately stored at -20°C for a maximum of one month before further processing.

For the *Ammodytidae* species mock community, specimens were caught during sampling efforts in 2021 and 2022 of *Ammodytes marinus* (55.5166 N, 03.7996 E)*, Ammodytes tobianus* (53.55718 N, 5.7346 E) and *Hyperoplus lanceolatus* (52.3790 N, 3.7997 E). All specimens were first morphologically identified and checked with a diagnostic primer pair. *H. lanceolatus* was further verified on 2 additional genes (Van Bleijswijk et al., *in prep.*). A fragment of ∼1mm^3^ tissue sample was taken.

### Primer design

Primer design is based on the adjacent ribosomal genes 12S and 16S of the mitochondrial genome of bony and cartilaginous fish present in the North Sea and further. Primer pair was designed *in silico* in Geneious prime 2019.0.4 (Kearse et al., 2012) and based on the in NCBI available mitochondrial genomes of the target species. A consensus sequence for each species was constructed when multiple genomes were available from the same species using default settings of the MAFFT alignment tool (v7.450, Katoh & Standley, 2013) incorporated in Geneious. Consensus sequences of all species were aligned and forward and reverse primers was designed manually by locating regions with low genetic variation between target species. This resulted in a long read universal fish primer pair (table 1) targeting a 2kb fragment from 450bp downstream the 12S rRNA gene in forward direction and 300bp upstream the 16S rRNA gene in reverse direction (figure 2A). The 5’ ends of the primers were extended with an ONT tag to allow for direct PCR based sample barcoding in downstream library preparation. To validate the 2kb primer pair, the primer pair was aligned against a curated North Sea database (see below) using Geneious prime 2023.0.4 (Kearse et al., 2012) in the “test with saved primers” mode (Primer3.2.3.7) allowing for 2 mismatches in the binding region. The primer pair was further validated with cutadapt v1.15 (Martin, Marcel, 2011) and showed that all mitochondrial sequences present in the database aligned with the primer pair in the correct region.

**Figure 2.**
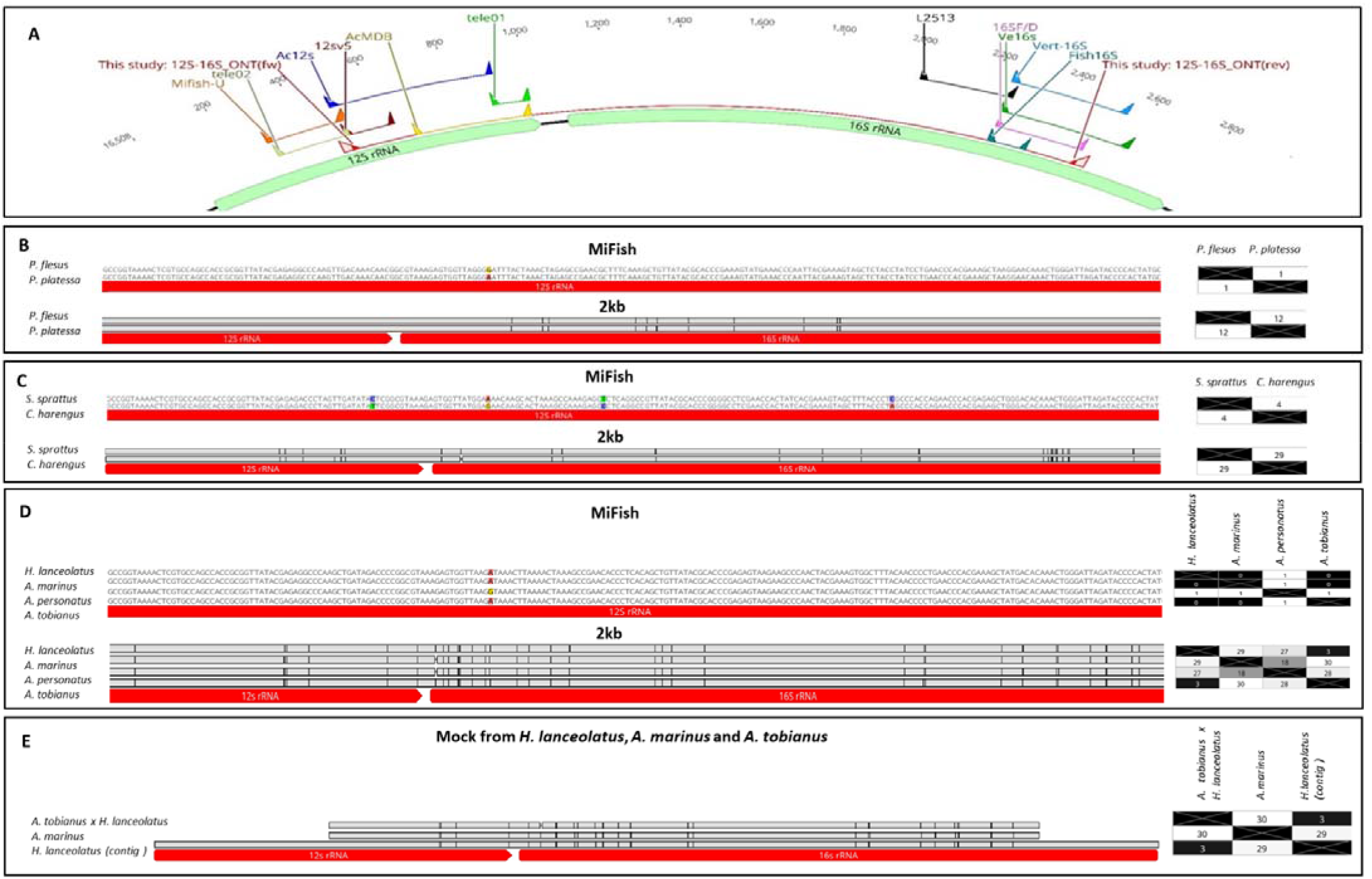
A) the position of the new 2kb primer pair (red) and among others the MiFish primer pair (orange). B) Genetic diversity between Platichtys flesus and Pleuronectes platessa for the target regions of the different primer pairs and C) S. sprattus and Clupea harengus and D) different Ammodytes species (Hyperoplus lanceolatus, A. tobianus, A. personatus and A. marinus). Matrices show the genetic dissimilarity (number of bp) between sequences within each Family(a, b, c). E) Experimentally measured genetic diversity of the 2kb fragment from amplicons of a mixed sample with three species of Ammodytes (H. lanceolatus, A. marinus and A. tobianus) and a contig generated from native sequencing of H. lanceolatus.

**Table 1:**
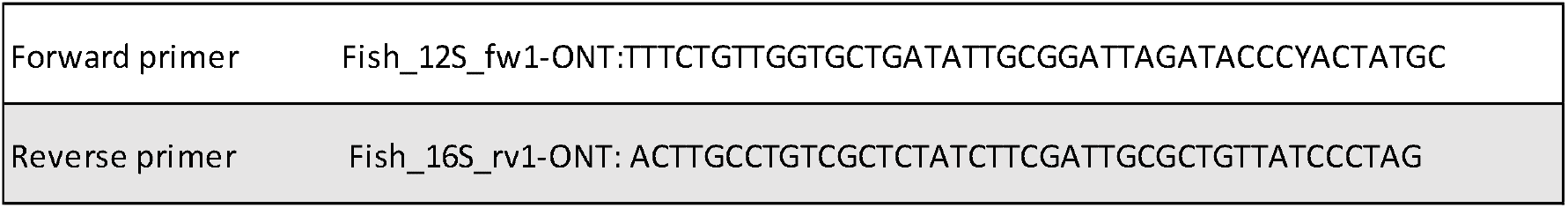
Primer sequence of the newly designed forward and reverse primer for the 2kb, including the ONT-specific primer extension enabling PCR barcoding (underlined).

### DNA extraction

To pre-process the samples, 20µL Proteinase K was added to the samples in DNA/RNA shield, together with 400µL lysis buffer of DNeasy Blood and Tissue extraction kit (Qiagen, USA) followed by 400µL 70% ethanol. Further DNA extraction was performed using this kit following the protocol for tissue samples. DNA concentrations were measured using a Qubit 2.0 Fluorometer (Invitrogen, USA). DNA from filters from the two North Sea datasets were extracted using the Quick-DNA miniprep kit (Zymo, USA) according to the manufacturer’s instructions. Details of both protocols are also given at protocols.io (dx.doi.org/10.17504/protocols.io.6yfhftn). Genomic DNA was isolated from *Ammodytidae* samples using the Gentra puregene DNA extraction kit (Qiagen, USA) according to manufacturer’s protocols. A *Ammodytidae* mock community was made by pooling these DNA extracts equimolarly.

### Amplification

For PCR amplification of the samples with the 2kb primer pair, 10µL 2x Phire Tissue Direct PCR Master Mix (ThermoFisher Scientific, USA) was used. To the master mix 0.4µL of each primer (10mM), 0.5µL eDNA template and nuclease free water (NFW) was added to a total of 20µL. Samples that were amplified with the MiFish primer pair consisted of 5µL 2x Phire Tissue Direct PCR Master Mix in combination with 1µL template and 0.2µL of each primer, and NFW added to a total of 10µL. To reduce the effect of stochastic heterogeneity in PCR amplification, each sample was amplified using 3 PCR replicates. For the amplification with the 2kb primer pair of the Ray Reef aquarium samples PCR settings were 98°C 180s, 98° 8s sec, 59.6°C for 10s, 72°C 30s, and 72°C 3min with 35 cycles and for the 2kb North Sea field samples the same amplification program was used, with 35 to 45 cycles depending on amplification yield (table S1). For amplification with the MiFish primer pair, PCR settings were: 98°C 180s, 98° 10 sec, 59.6°C for 8, 72°C 10s, and 72°C 3min with 35 cycles. PCR replicates were pooled prior to purification with magnetic beads.

### Nanopore sequencing

For the North Sea field samples amplified with the 2kb primers (both Borkum Reef/Gemini Wind Park transect and wrecks) amplicons were barcoded using the PCR barcoding kit 96 (EXP-PCB096), and the sequence library was created using the SQK-LSK109 kit. All other samples were also barcoded using the PCR barcoding kit 96 (EXP-PCB096), and sequencing libraries were created using the SQK-LSK114 kit (Oxford Nanopore Technologies Ltd., UK). The following adaptations were made from the manufacturer’s instructions: barcoding PCR was performed in a total volume of 15µL containing 0.3µL PCR barcode primer pair and 10-50ng amplicon. The applied barcode PCR program was as follows: initial denaturation at 95°C for 180s, 15 cycles of 95°C for 15s, 62°C for 15s (10s for MiFish), 65°C for 90s, followed by a final extension at 65°C for 180s. After the barcoding PCR, sample concentration was estimated using the Qubit HS kit on the non-purified barcoded PCR products, and samples were pooled in equimolar ratios. The pooled amplicon sequence library was cleaned using magnetic beads, washed once with 70% ethanol and once with a 2:1 mixture of Long Fragment Buffer (LFB) and Short Fragment Buffer (SFB) (LFB and SFB are supplied with the Ligation sequencing kits from ONT) to enrich for the 2kb target size fragments, and only SFB for MiFish samples. During final clean-up, the library was again washed in a 2:1 mixture of LFB and SFB (2kb) or SFB only (MiFish). A maximum of 100ng DNA was loaded on a primed flow cell to prevent overloading of the flow cell. If necessary, in case of the SQK-LSK109 library, the R9.4.1 flow cell was refuelled using a mixture of Sequencing Buffer (SQB) and nuclease free water (1:1). For the 2kb samples, sequencing was performed until an average sequencing depth of 10-50k reads was obtained per barcode for the Borkum Reef/Gemini Wind Park samples and 70-100K per barcode for the shipwreck samples. MiFish samples were sequenced with an average sequencing depth between 50-300k reads per samples. Sequencing of the 2kb North Sea field samples was performed using an Oxford Nanopore Mk1b in combination with an R9.4.1 flow cell, whereas the Aquarium 2kb, *Ammodytidae* samples and MiFish samples were sequenced on a Mk1C with an R10.4.1 flow cell with a sequencing speed of 450 bases per second. Different sequencing chemistries were combined in this study due to the quick upgrades of the Nanopore chemistries and therefore the R9.4.1 flow cells orders were discontinued within the lab during additional analysis of the samples.

### Sequence read processing

Base-calling of the raw fast5 files was performed using Guppy (Version 6.5.7, Oxford Nanopore Technologies Ltd., UK) in super high accuracy (SUP) mode. For the bioinformatic analysis, Nanofilt v2.3.0 was used to filter raw reads on quality and read length (De Coster et al., 2018). Then cutadapt v1.15 (Martin, 2011) was used to trim the extensions and primers (CD-hit v4.8.1), a program that clusters reads based on short words rather than sequence alignment was used to cluster the reads based on a set percentage of similarity (W. Li et al., 2002) The clustered reads are subsequently aligned using Minimap2 v2.17 (H. Li, 2018). Based on these alignments, Racon v1.4.13 is used to build the initial draft consensus sequence of each cluster (Vaser, Sović, et al., 2017) which is then polished by Medaka v1.1.2 (Oxford Nanopore Technologies Ltd., UK). We compiled a pipeline, called Decona, to automate these processing steps (https://github.com/Saskia-Oosterbroek/decona). The raw basecalled reads are first trimmed with a read length filter at 1800-2350 bases for fragments of the 2kb primer pair dataset and between 150 and 240 for fragments of the MiFish primer pair dataset. A cluster similarity at 85% was chosen based on carefully checking different clustering percentages (data not shown). Medaka polished consensus sequences were created from each cluster larger than 5 reads. Initial polished consensus sequences were re-clustered at 99%, and when the cluster size exceed 500 reads, clusters were subsampled to a maximum cluster size of 500 reads. The commands used to run Decona were as follows:(2kb: decona -f -T 20 -l 1800 -m 2350 -g “GGATTACCCYACTATGC;max_error_rate=0.1;min_overlap=17…CTAGGGATAACAGCGCAATC; max_error_rate=0.1;min_overlap=17” -n 5 -r 0.99= -R 500 -M) (MiFish: decona -f – T 20 -l 160 -m 240 -g “GTYGGTAAAWCTCGTGCCAGC;max_error_rate=0.1;min_overlap=20…CAAACTYGGATACCCCACTAT; max_error_rate=0.1;min_overlap=20” -n 5 -r 0.99= -R 500 -k 6 -M).

### Curated North Sea fish reference database building

For taxonomic identification, an in-house reference database was compiled based on whole mitochondrial genome sequences available in the NCBI database for North Sea fish species (last search April 2023). When the whole mitochondrial genome was not available, available sequences of (fragments of) the 12S and/or the 16S rRNA genes from these species were added to the database. To validate correct species identification, closely related species that do not occur in the North Sea were also added to the database as were frequently occurring contaminants. The complete database consisted of 532 sequences of which 111 were complete mitogenomes and 30 were complete 12S and 16S regions (table S3). The database contained 224 unique species.

### Taxonomic assignment

Species identifications of the 2kb consensus sequences were obtained with the taxonomic identifier Centrifuge v1.0.4 (Kim et al., 2016) with a minimal alignment length of 200 nucleotides and using only 1 primary assignment for each consensus sequence. If a consensus sequence was aligned with the same quality/score to two or more species, sequences were assigned to genus level and not further processed. Consensus sequences that could not be identified on species level were labelled unclassified. A minimal alignment length of 200 nucleotides gave the highest species richness that still yielded meaningful results, meaning North Sea vertebrate hits. In addition, 2kb sequences were also classified using a BLASTn search (NCBI, version 2.11.0) against our North Sea fish reference database. Top hits were considered accurate on species level when there was a minimal alignment length of 500 nucleotides with <15 mismatches and >99% identity. With percentage identities >98% sequences were assigned on genus level and with >97% sequences were assigned at family level but not further considered for this study. Due to the use of the longer 2kb sequences, Centrifuge considerably outperformed BLAST in terms of overall correct species identification (table S4) but BLAST also identified a large proportion of the sequences that could not be assigned by centrifuge. Therefore it was decided that the results from both datasets were merged. The MiFish sequences were classified with BLASTn using our North Sea fish reference database. Species level assignment was accepted when the top hit had <4 mismatches, >99% identity, and a minimal alignment length of 100 nucleotides to exclude potential chimeric reads.

### In sillico comparisons of species groups with little interspecific genetic differences

*In sillico* comparative alignments were made from different taxonomic groups relevant for this study (e.g sharks, rays, wrasses, gurnards, flatfishes, gobies, sand eels, mullets etc, data not shown) of (partial) mitochondrial references from the NCBI database. Genetically closely related species were aligned using Muscle 5.1 (Edgar, 2004) multiple alignment tool in Geneious prime (table S5, for accession numbers). Target regions of the different primer pairs were identified using saved primers mode (Primer3.2.3.7) allowing for 2 mismatches in the binding region.

### Native sequencing and analysis of Hyperoplus lanceolatus mitogenome

Besides amplicon sequencing total genomic DNA isolated from *H. lanceolatus* tissue was separately sequenced using SQK-LSK114 kit (Oxford Nanopore Technologies Ltd., UK), according to the manufacturer’s instructions, using a MinION MK1C and a R10.4.1 flow cell, and a sequencing speed of 450 bases per second. We sequenced 2.48gbase of reads, with an N50 of 553bp. Reads were basecalled using Guppy (Version 6.5.7, Oxford Nanopore Technologies Ltd., UK) in super high accuracy (SUP) mode. Reads were analysed using a protocol to assemble and annotate Nanopore based mitochondrial genomes (Oosterbroek et al., *in prep*).

### Analysis of taxonomic assignments

The taxonomically assigned output data were further analysed in R studio (2022.12.0). Taxonomic lineage was obtained from NCBI using taxize v0.9.96 (Chamberlain & Szöcs, 2013). As tag leakage is observed within the 10.4.1 chemistry of ONT, a tag leakage correction was performed. For this correction, we calculated the total read number for each taxon from the complete sequencing run. Next, 0.1% from this taxon specific number was subtracted from each individual barcode to correct proportionally for the total read abundance of each taxon. As such, all species that had a total read occurrence of more than 1000 reads in the total dataset received a correction in the barcode specific results. Species accumulation curves were plotted (vegan package, v 2.6-4, table S6) and the sequence abundance was rarefied to 50,000 reads per sample (MiFish) 10,000 reads per sample (Aquarium and *Ammodytidae*) and 5000 reads per sample (North Sea field samples) using phyloseq v1.30.0 (McMurdie & Holmes, 2013). Reads classified as belonging to the genera *Homo*, *Ovis*, *Gallus*, *Bos* and other non-marine animals were set to unclassified, along with all consensus sequence that did not have a hit with a database on species level (see table S7 for the read percentage of non-fish hits per barcode). Control samples were analysed but not considered for further analysis (table S8). For alpha diversity, both Shannon indices and observed values were calculated and were tested using Shapiro-Wilk for normal distribution of the data, one or two-way ANOVA to test for significant differences between alpha diversities and post hoc Tuckey HSD test for pairwise comparison. For beta diversity, non-metric multidimensional scaling (‘bray’) was performed in combination with betadisper to check for homogeneity of variance and PERMANOVA to analyse the effect of treatments between samples (vegan). Post-hoc analysis was performed using the pairwise.adonis package in combination with devtools.

## Results

### *In sillico* and *in vitro* comparison of primer pair performance on closely related taxa

Although our primer design was based on mitogenomes of bony fish and elasmobranch only, the resulting primers turned out to be universal not only to fish, but also to almost all other vertebrates. As a result, we regularly pick up not only marine mammals and birds, but also contaminating DNA of human, chicken, cow and pig. From the *in sillico* alignments, *Pleuronectes platessa* and *Platichthys flesus* target regions differ 1 nucleotide the MiFish primer pair (99.4% similarity) whereas the target region of the 2kb primer pair has 12 nucleotide differences (99.3% similarity) (figure 2B). *Clupea harengus* and *Sprattus sprattus* diverged by 4 nucleotides (98.3% similarity) in the MiFish target region and their 2kb target region showed a pattern of 29 nucleotide differences (98.6%) (figure 2C). Sand eel species *Ammodytes marinus*, *A. tobianus* and *H. lanceolatus* and *A. personatus* also show 1 nucleotide mismatch (99.6% similarity) between all species for the MiFish target region. From the 2kb target region, *A. marinus* differed from *A. tobianus* and *H. lanceolatus* with 29 and 30 nucleotide differences respectively, (98,3% similarity) whereas between *A. tobianus* and *H. lanceolatus* the genetic diversity remains low with 3 nucleotide differences (99,1%) (figure 2D).

Next to the *in sillico* comparison we also tested the combination of amplification of the 2kb primer pair target region, Nanopore amplicon sequencing and Decona data processing on a mixed sample of genomic DNA extracted from three species of sand eels. BLASTn top hits obtained of consensus sequence show a separation between *A. marinus* and *A. tobianus*, whereas no top hit match with *H. lanceolatus* was found. Alignment of reference sequences for the 2kb target region verified the high similarity between *H. lanceolatus* and *A. tobianus* with a nucleotide difference of 3 (99.1%) (figure 2E). To validate this, we separately sequenced total genomic DNA of *H. lanceolatus* without amplification, and assembled a contig spanning the 2kb region of the *H. lanceolatus* mitochondrial genome (GenBank: OR209138), indeed demonstrating that the sequence identity between these two species differs only by 3 nucleotides for that region. Outside of the 2kb region more nucleotide differences were detected (not shown). The absence of consensus sequences showing the *H. lanceolatus* version is likely due to clustering and polishing steps. These were able to correctly separate *A. tobianus* and *A. marinus*, but removed the variation between *A. tobianus* and *H. lanceolatus.*

### Enhanced assignment accuracy up to 100% using Decona

Of all consensus sequences generated by Decona that were identified with BLASTn, 33% could be correctly assigned based on a percentage identity of 99% or higher. Of these, 42% of the 2kb fragment consensus sequences and 82% of the MiFish consensus sequences had a BLASTn top hit to the reference database with a percentage identity of 99.9% or 100%.

### Similar alpha diversity obtained with both primer pairs in ray reef aquarium, but species composition varies

From the North Sea Ray Reef aquarium, 221995 (2kb) and 220963 (MiFish) basecalled reads were obtained from the 2kb approach, of which 50949 (2kb) and 152395 (MiFish) reads with a barcode distribution of 25474±13483 (2kb) and 76198±8010 (MiFish) reads per barcode were used for consensus building (table S2). Shapiro-Wilk showed normally distributed data (Shannon: P=0.818, Observed P=0.415) and no significant difference in evenness (Shannon, ANOVA p= 0.824, figure 3A) or richness (observed, ANOVA P=0.415, figure 3A).

**Figure 3.**
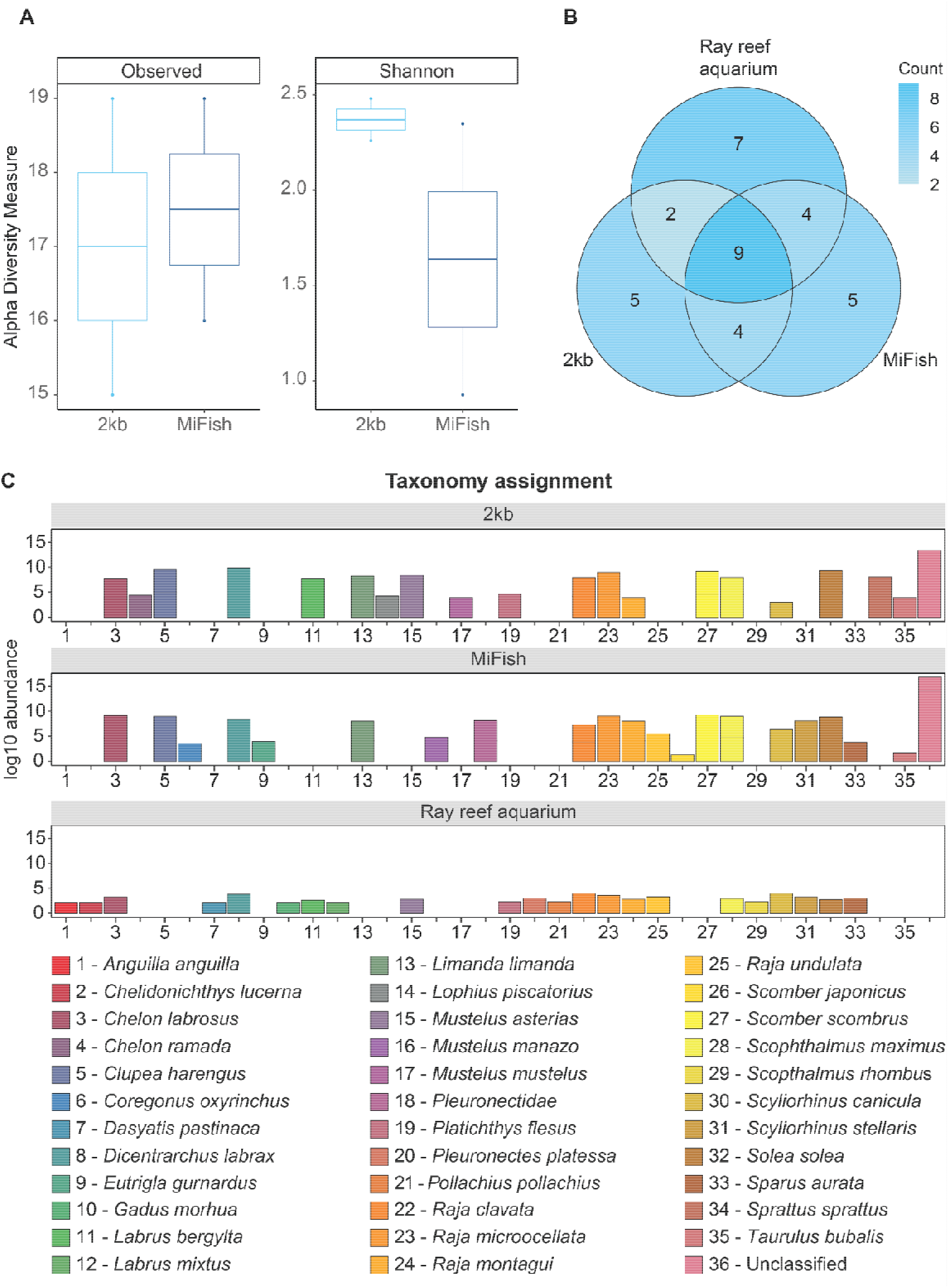
MiFish and 2kb primer pair in comparison to the species composition of the North Sea Ray aquarium. Taxonomy assignment on species level. A) Alpha diversity comparison of richness and evenness, B) Venn diagram showing the species overlap in morphological species counts and the primer used C) Barplot of the species specific differences observed between the actual species present in the Ray aquarium and the results from the analysis using the two different primer pairs.

Of the 22 North Sea vertebrates present in the aquarium, a total of 9 species could be identified with both primer pairs of which an additional 2 species could be obtained with 2kb sequence assignments (figure 3B). An additional 4 species could be identified with MiFish sequence assignments. Four species were identified with both primer pairs but were not reported as aquarium inhabitants. Both primer pairs amplified fragments that were assigned to 5 species not identified with the other primer set. Of these species, sequences from both primer pairs were incorrectly assigned to species that belonged to a genus of which the correct species was also taxonomically assigned (figure 3C). A total of 7 species that resided in the aquarium could not be detected in the eDNA samples by either of the primer pairs. Of these, 5 species had only one individual in the aquarium. For the two undetected species that did have multiple individuals in the Ray reef aquarium, *Scopthalmus rhombus* was not present in the reference database of either primer pair.

### Significant difference in alpha and beta diversity in North Sea field samples from different habitats

From the North Sea samples of Borkum, Gemini and sandy bottom, 235467 (2kb) and 1620934 (MiFish) basecalled reads were obtained from the 2kb approach, of which 95217 (2kb) and 981304 (MiFish) reads with a distribution of 9992±5907 (2kb) and 108864±88918 (MiFish) reads per sample were used for clustering and consensus building (table S2). For alpha diversity, samples were normally distributed (Shapiro-Wilk, Shannon p=0.199, Observed p=0.132) and a significant difference of richness was found between primer pair choices (Observed, 2-way ANOVA, p= 0.0003) and location (Observed, 2-way ANOVA, p= 0.003) (figure 4A). Also a significant difference in evenness was found between primer pair choices (Shannon 2-way ANOVA, p= 0.009) but not between location (Shannon, 2-way ANOVA, p= 0.78). No significant interaction effect was found between alpha diversity measures (Observed, p=0.616, Shannon, p=0.213) (figure 4A). For the beta diversity the NMDS ordination plot shows clustering of sampling triplos within each location (indicated in colours) and within each location, also clustering between primer pair choice can be observed (indicated in shapes) (figure 4B). These effects of location and primer pair choice were verified with statistical analysis. Homogeneity of variances between samples was found (betadisper p=0.539) and PERMANOVA showed a significant effect of primer pair choice (adonis, p=0.001) and location (adonis, p= 0.001) and no significant interaction (adonis p=0.106). Pairwise analysis furthermore found a significant difference between Gemini and the Borkum reef (pairwise.adonis, p= 0.009) and the Borkum reef and sandy bottom (pairwise.adonis, p=0.006) but not between Gemini and Sandy bottom (pairwise.adonis, p=0.492). More statistics calculation details are given in table S9. Both primer choices showed that unique species were observed with either method. *Limanda aspera*, *Gadus morhua* and *Trisopterus luscus* were only observed using the MiFish primer pair whereas *Ammodytes marinus*, *Limanda limanda*, *Taurulus bubalis*, *Raja clavata* and *Callyonimus lyra* are unique for the 2kb primer. pair (figure 4C).

**Figure 4.**
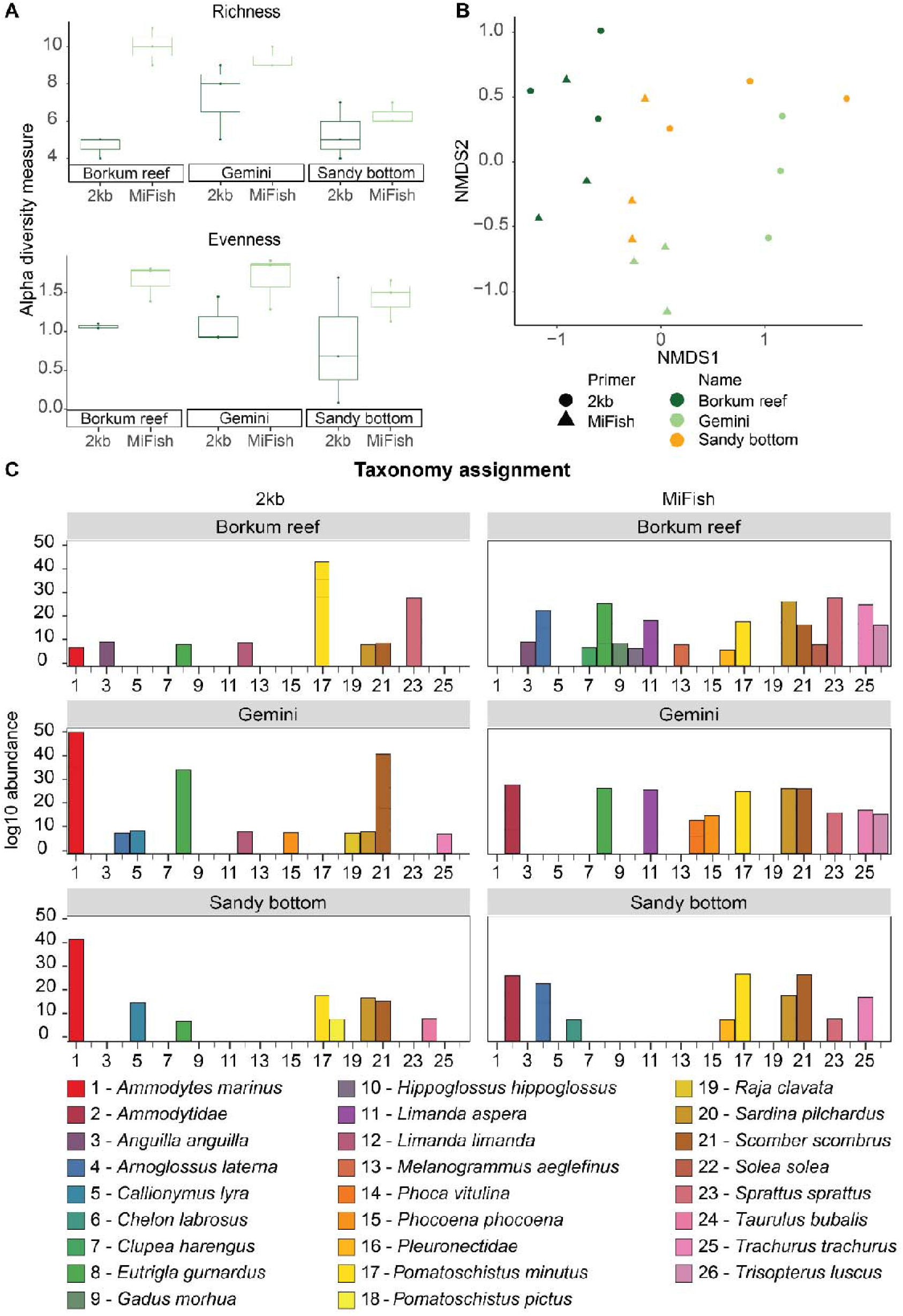
MiFish and 2kb primer pairs comparison of different eDNA samples taken in the Borkum reef, Gemini wind park and Sandy bottom. Taxonomy assignment on species level. A) Alpha diversity comparison of richness and evenness, B) NMDS ordination plot (Bray) showing the similarity between samples C) species comparison barplot of the species specific differences between the different location and primers.

### eDNA samples taken at different ship wrecks significantly differ in alpha and beta diversity

From samples of the ship wreck, 228015 basecalled reads were obtained, of which 111598 reads with a barcode distribution of 12022±2399 reads per barcode were used for clustering and consensus building (table S2). Shapiro-Wilk showed normally distributed data (Shannon, p=0.09, Observed p=0.306) and no significant difference in richness (Observed, one way ANOVA p= 0.131, figure 5A) but a significant difference in evenness (Shannon, one way ANOVA p=0.009) was observed (figure 5A). There was a significant difference between wreck 1 and 3 (p=0.024) and 2 and 3 (p=0.011) but not between wreck 1 and 2 (p=0.745). For beta diversity, the NMDS ordination plot shows clustering between each wreck (indicated in colors) (figure 5B) and PERMANOVA showed a significant difference in beta diversity between wrecks (adonis, p=0.001) and homogeneous samples (betasisper, p=0.106). Species composition indeed showed location specific community compositions (figure 5C). Interestingly we have also found assignments of the marine mammal *Phocoena phocoena* in shipwreck 1.

**Figure 5.**
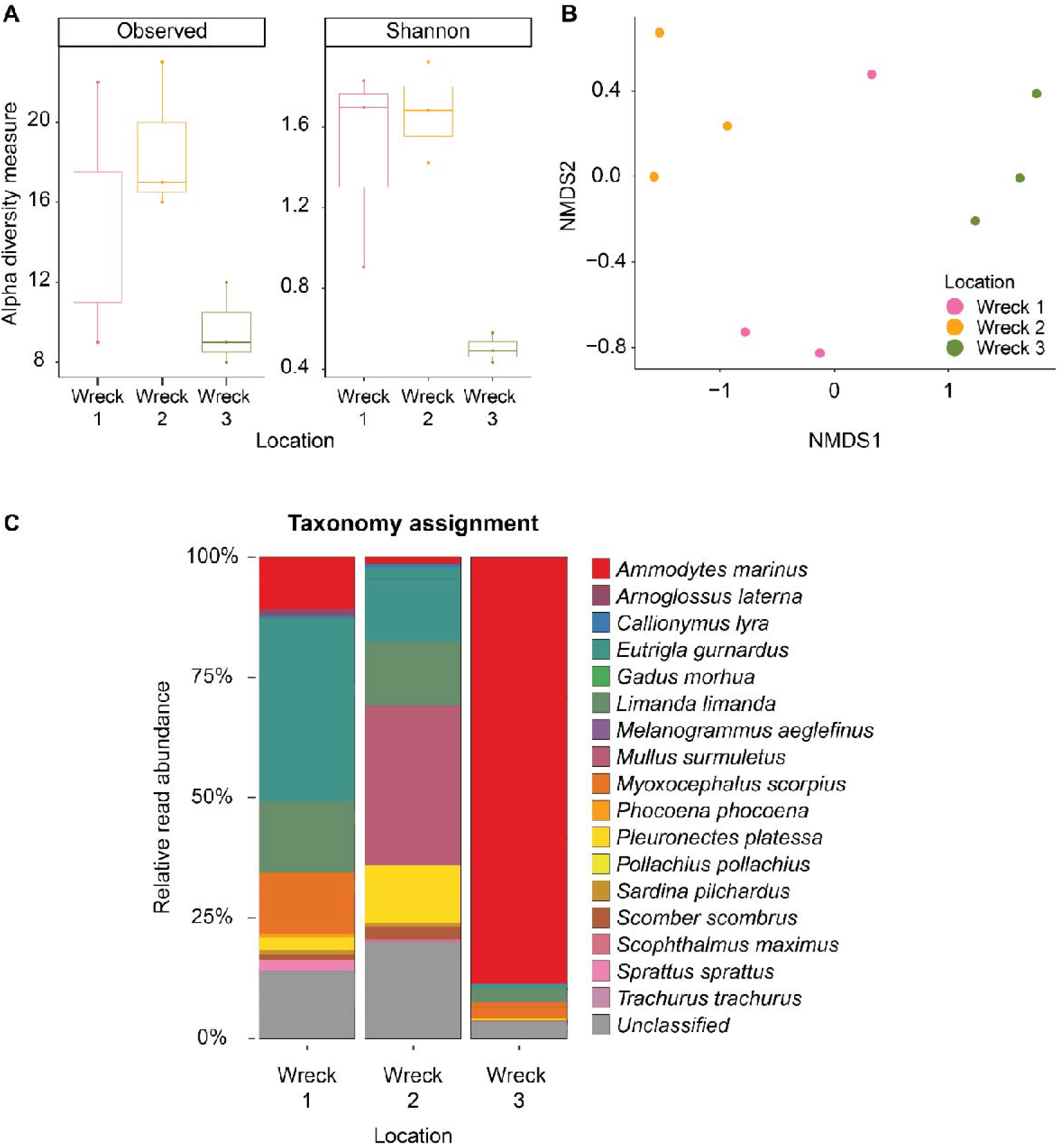
Relative read abundance of marine vertebrate species assignments associated with different ship wrecks. A) Alpha diversity comparison of richness and evenness, B) NMDS ordination plot of the beta diversity (Bray) C) barplot of the species specific differences between the wrecks.

## Discussion

eDNA based metabarcoding has revolutionized taxonomic identification and monitoring of marine ecosystems (Thomsen & Willerslev, 2015). This resulted in a steep increase in usable molecular markers (Deagle et al., 2009; Devloo-Delva et al., 2019; Leray et al., 2013; Miya et al., 2015b), paralleled with the fast increase of genetic databases for marine organisms (Benson et al., 2013; Federhen, 2012; Manel et al., 2020; Ratnasingham & Hebert, 2007; Schroeter et al., 2020). With the rise of Nanopore sequencing, it now becomes increasingly feasible to use long read sequencing in metabarcoding studies. Although ONT based long read metabarcoding is shown to be successful in bacterial studies (Krehenwinkel et al., 2019; Matsuo et al., 2021), it has thus far not been developed and validated for marine biodiversity studies.

We designed a primer pair for marine fish that turned out to be specifically targeting all marine (and terrestrial) vertebrates. The 2kb fragment spans both 12S and 16S regions, often used in metabarcoding approaches. We compared this new primer pair with the commonly used universal short fragment MiFish primer pair. We first analyzed the potential sensitivity and specificity in an *in sillico* approach, and subsequently tested them both on samples with a known composition and finally on field samples. In doing so, we have demonstrated the benefit of using long read sequencing in marine eDNA samples by using Nanopore sequencing. Adequately demonstrating the possible strength of long read amplicon sequencing remains challenging, as experimental and methodological choices are often different between studies (Ruppert et al., 2019b; Wang et al., 2021). Also in this study we do show an limited amount of samples, as offshore marine conditions often restrict the amount of samples that can be effectively taken. In addition, we did not include the comparison with a different sequencing platform such as the Illumina MiSeq system, since several authors already reported the comparability between both sequencing platforms (Heikema et al., 2020; Van Der Reis et al., 2023). Our comparative study was focused on demonstrating the added benefits of analyzing longer amplicon fragments over commonly used short amplicons, and helps building a fundament for long read sequencing in marine vertebrate monitoring approaches.

### Enhanced accuracy up to 100% using Decona

The developed sequence read processing pipeline Decona enables increased consensus sequence accuracy up to 10% points compared to the basecalled read, comparable to Illumina read accuracy (Caporaso et al., 2011). There is limited automated bioinformatics processing reported in Nanopore based studies, especially for metabarcoding (Santos et al., 2020b). This study contributes by making the Decona pipeline available (https://github.com/saskia-oosterbroek/decona). Once installed, one line of code suffices to correctly run the pipeline and enables data processing for scientists with limited experience in the command line. The bioinformatics tools integrated in Decona are well established programs in genomics and transcriptomic studies. For example, tools as CD-Hit (Huang et al., 2010) have previously been used in analysis of Nanopore sequence data for clustering and consensus building of fish amplicon based sequences (Voorhuijzen-Harink et al., 2019). Reference based polishing was successfully applied when identifying benthic organisms on autonomous reef monitoring structures (Jin et al., 2020) using *minibarcoder.py* (Srivathsan et al., 2018). The combination of both clustering and *de novo* alignment based polishing with racon (Vaser, Sović, et al., 2017) and medaka (https://nanoporetech.github.io/medaka/) has previously been used for the correction of metagenomes (Rodríguez-Pérez et al., 2020). In contrast, the Decona pipeline combines similarity based clustering based on short word tables instead of an alignment approach in combination with alignment based polishing with racon and medaka, which further increases the sequence accuracies. Limitations of Decona may lay in the necessity to cluster, which makes it very well possible that reads from genetically similar organisms end up in the same clusters, resulting in lower detection sensitivity then is actually sequenced. In addition, clustering with Decona also disregards singletons and reads and clusters consisting of less than 5 reads, as such missing the rare reads in datasets. Fortunately, due to the fast development of Oxford Nanopore sequencing technologies, new sequencing chemistries and base calling algorithms are often released and accuracy is now at a 99.8% raw read accuracy for model organisms (Srivathsan et al., 2021). It becomes therefore possible to skip the clustering and polishing process al together, and perform raw read identification using amplicon sequence variants (ASVs) with the new chemistries, as is the standard for Illumina platforms (Van Der Reis et al., 2023).

### In sillico identification shows increased species level identification using longer DNA fragments

Alignment of species within the *Ammodytidae, Pleuronectidae* and *Clupeidae* families all showed a genetic variability insufficient to differentiate related species when aligning the MiFish target fragments. These assignment problems have already been reported for North Sea fish species (e.g. Barco et al., 2022). The 2kb target fragment alignment shows that for some species indeed taxonomic variability increases to a dissimilarity up to 2%. However, for *H. lanceolatus* and *A. tobianus,* it remains impossible to distinguish on species level on the complete 2kb target region, which demonstrates that for some species it is needed to use an additional target region to adequately assign on species level. In addition, for *P. platessa* and *P. flesus* there is an increased amount of nucleotide differences found on the 2kb target region, however this difference is still below the chosen threshold for species level identification in this paper. This *in silico* comparison therefore shows that enhanced identification resolution on species level can be obtained when using long read metabarcoding, but that there is a need to consider other fragments and lengths to be able to differentiate between genetically highly similar species, or, alternatively, accept lower thresholds when comparing longer fragments.

### Assignments from the North Sea Ray Reef aquarium show overlap between primer pair choices, primer pair specific differences, and false positives

In the North Sea Ray Reef aquarium, the majority of the species could be identified with both primer pairs used. The species *S. sprattus* and *S. scombrus* were identified by both primer pairs but did not live in the aquarium. These species were used as feed for the different piscivores animals in the aquaria, showing that both primer pairs are also able to pick up the signal of the diet of these animals (personal communication Paulien Bunskoek, Dolfinarium Harderwijk).

Analysis of the eDNA samples with the primer pairs described here also shows the detection of unique species for each primer pair (figure 5), for example the sequences from the MiFish primer pair better represent the different ray species. This results agrees with earlier findings illustrating that multiple markers give a better representation of the complete biodiversity (Cordier et al., 2019). For both primer pairs there is also a certain degree of false positive species assignment of the sequences observed as is exemplified with *Chelon ramada* and *Mustelus mustelus*. For longer sequences, the genomic identifier Centrifuge is used with an alignment length of 200bp. This was initially chosen because it gives the highest amount of meaningful species. However, such threshold may also result in false positives, because identification tools such as Centrifuge and BLASTn will assign higher scores to longer alignments. This can cause a bias in genera where the longest matching reference sequence in the database is that of that of a related species instead of the actual target species.

There were also species present in the aquarium that were not identified by any primer pair. It is often observed that eDNA methods do not identify the complete biodiversity, despite using a multi marker approach (Morey et al., 2020). *S. rhombus* for example was not identified by any molecular method as there is no representation of its 12S and or 16S rRNA gene in public databases, making it impossible to align to this species. This further stresses the need to continue improving genetic reference databases both with short fragments as well as for complete (mitochondrial) genomes. Of the other undetected species only one individual per species was present in the aquarium, which suggests that the lack of detection is a result of low initial DNA concentration. This is in line with inconsistent detection of rare taxa between filters described in previous reports (Evans et al., 2017; Kelly et al., 2019; Morey et al., 2020), and species detection could be improved by using more replicates (Beentjes et al., 2019; Evans et al., 2017) or a larger volume of water where possible. An alternative explanation for the lack of detection of these low abundant species could come from the sequence processing. As it is necessary to cluster raw reads, rare reads can end up in a cluster that is removed during further sequence processing, as Decona discards clusters smaller than 5 reads. Overall, despite the detection of false positives, false negatives and primer pair specific results, both primer pairs could identify the majority of the marine vertebrates and there was no qualitative difference between primer pair choices.

### Field samples show similar beta diversity between different North Sea habitats

The alpha diversity for both evenness and richness was higher for the MiFish results in the North Sea and is in contrast with non-significant differences in alpha diversity between used primer pairs in the Ray reef aquarium. An explanation for this increase in alpha diversity could lie in the eDNA fragment length sampled. Opposed to aquaria where the relative concentration of eDNA in the water and especially of freshly released eDNA in the water is higher, field samples have a lower fish concentration per liter of water, and potentially faster breakdown of free extracellular DNA (Seymour et al., 2018). Therefore, it is likely that there are fewer intact DNA fragments present in the field eDNA samples that can be successfully amplified with the 2kb primers, whereas amplification of short DNA fragments with the MiFish primer pairs might still be possible. This suggests that the MiFish primer pair approach can identify additional signals from taxa that released their eDNA longer ago. Despite this difference, similar patterns of beta diversity for both primer pairs have been observed, showing that both primer pairs can describe habitat dissimilarity equally well. This difference could be exploited by using short read primer pairs in combination with long read primer pairs to obtain additional insight into the diversity on a temporal scale.

At two sample locations, *A. marinus* could be identified by the 2kb primer pair, while this species level identification could not be made using the MiFish approach (figure 3). *Limanda* sequences from the MiFish primer pair also consistently were assigned to *Limanda aspera*, which is the Chinese yellowfin sole that does not occur in the North Sea. Similar observations of false positive identifications were seen in the Ray reef aquarium with *S. japonicus* instead *S. scombrus* (figure 4). As the BLASTn function was used for taxonomic assignment of sequences from the MiFish primer pair, it is possible that the top hit passed quality control for the wrongly assigned species while there was also a similar match with the correct species. Taxonomy assigners that are currently used for Illumina MiFish metabarcoding make use of Naïve Bayesian classifiers such as RDP (Cole et al., 2003) that is able to quickly do taxonomic assignment for ASV metabarcoding sequences and also assigns taxa to a higher taxonomic level when there are sequences with multiple hits. For Nanopore based consensus sequences such assigners are not yet applied, which results in manual adjustments of results and requires a priori knowledge on the genetic similarity between species to detect such false positive results.

We demonstrated that application of the 2kb primer pair on eDNA field samples is sensitive enough to find spatial differences between sampling sites. The monitoring of fish biodiversity at multiple different sample sites (e.g. ship wreck sampling) gives a good overview of the vertebrate richness on a local scale and additionally provides insights into visiting vertebrate species such as the harbor porpoise.

## Conclusion

This study demonstrates and validates an eDNA metabarcoding approach using Nanopore long read technology, enabling increased resolution on species level. We demonstrate an increased species resolution due to the longer DNA fragments analyzed, enhanced by our Nanopore raw read processing pipeline Decona. We further show limitations such as false positive assignments and limited detection of rare species. This study also shows the importance of using multiple markers for increased detection resolution for fish and suggests to explore the possibility of using long read metabarcoding to gain biodiversity information on a spatio-temporal scale. Further research should focus on the use of Nanopore generated raw reads, and further implementing Nanopore based (long read) metabarcoding as standard to the molecular ecology toolbox. Moreover, it is essential that addition of longer reference sequences to databases, preferably of full (mitochondrial)genomes, maintains a high priority in marine molecular ecology. Only then long read based DNA metabarcoding and metagenomics can develop to its full potential to serve as monitoring tool.

## Acknowledgements

We are grateful for the members of ‘Duik the Noordzee Schoon’ foundation for providing the trip to the shipwrecks and assisting during the dives and sampling. We would like to thank the Dolfinarium in Hardewijk for letting us sample in their Ray reef aquarium and providing species information and metadata of the aquarium. We would like to thank Linda Tonk of Wageningen Marine Research, Miriam Schutter of Bureau Waardenburg and crew members of MS Vrijheid III, for providing the cruise to the Borkum Reef and Gemini Wind Park and assistance with sampling. We acknowledge GEANS, an Interreg project supported by the North Sea Programme of the European Regional Development Fund of the European Union and JIP ECO-FRIEND (RVO reference number TEWZ118017) for funding parts of this research. We thank Judith van Bleijswijk for supplying the Ammodytes specimens. Especially valuable contributions are made by Aline Joustra by providing the illustrations of the experimental design (http://www.alinesci.com/).

## Authors contributions

KD LJ and RN designed the experiment; LJ, PK, OB, RN, AM and EW were involved in sample collection and processing; KD, LJ and RN did the laboratory work; SO, KD and RN designed the bioinformatics pipeline Decona; KD and RN conducted the data analysis; KD, LJ, SO, RN and AM interpreted the data; all authors wrote and revised the manuscript.

## Data availability

All sequence data used for the writing of this manuscript have been uploaded to the NCBI SRA database, SRA submission number: SUB13560265.

## References

1. Andersen, J. H., Al-Hamdani, Z., Harvey, E. T., Kallenbach, E., Murray, C., & Stock, A. (2020). Relative impacts of multiple human stressors in estuaries and coastal waters in the North Sea–Baltic Sea transition zone. Science of the Total Environment, 704, 135316. https://doi.org/10.1016/j.scitotenv.2019.135316

2. Balasingham, K. D., Walter, R. P., Mandrak, N. E., & Heath, D. D. (2018). Environmental DNA detection of rare and invasive fish species in two Great Lakes tributaries. Molecular Ecology, 27(1), 112–127. https://doi.org/10.1111/mec.14395

3. Baloğlu, B., Chen, Z., Elbrecht, V., Braukmann, T., MacDonald, S., & Baloglu, B. (2020). A workflow for accurate metabarcoding using nanopore MinION sequencing 1. https://doi.org/10.1101/2020.05.21.108852

4. Barco, A., Kullmann, B., Knebelsberger, T., Sarrazin, V., Kuhs, V., Kreutle, A., Pusch, C., & Thiel, R. (2022). Detection of fish species from Marine Protected Areas of the North Sea using environmental DNA. Journal of Fish Biology. https://doi.org/10.1111/jfb.15111

5. Beentjes, K. K., Speksnijder, A. G. C. L., Schilthuizen, M., Hoogeveen, M., & Van Der Hoorn, B. B. (2019). The effects of spatial and temporal replicate sampling on eDNA metabarcoding. PeerJ, 2019(7), e7335. https://doi.org/10.7717/peerj.7335

6. Beng, K. C., & Corlett, R. T. (2020). Applications of environmental DNA (eDNA) in ecology and conservation: opportunities, challenges and prospects. In Biodiversity and Conservation (Vol. 29, Issue 7, pp. 2089–2121). https://doi.org/10.1007/s10531-020-01980-0

7. Benson, D. A., Cavanaugh, M., Clark, K., Karsch-Mizrachi, I., Lipman, D. J., Ostell, J., & Sayers, E. W. (2013). GenBank. Nucleic Acids Research, 41(D1), D36–D42. https://doi.org/10.1093/nar/gks1195

8. Bessey, C., Jarman, S. N., Berry, O., Olsen, Y. S., Bunce, M., Simpson, T., Power, M., McLaughlin, J., Edgar, G. J., & Keesing, J. (2020). Maximizing fish detection with eDNA metabarcoding. Environmental DNA, edn3.74. https://doi.org/10.1002/edn3.74

9. Bleidorn, C. (2016). Third generation sequencing: Technology and its potential impact on evolutionary biodiversity research. In Systematics and Biodiversity (Vol. 14, Issue 1, pp. 1–8). Taylor and Francis Ltd. https://doi.org/10.1080/14772000.2015.1099575

10. Bleijswijk, J. D. L., Engelmann, J. C., Klunder, L., Witte, H. J., Witte, J. IJ., & Veer, H. W. (2020). Analysis of a coastal North Sea fish community: Comparison of aquatic environmental DNA concentrations to fish catches. Environmental DNA, edn3.67. https://doi.org/10.1002/edn3.67

11. Caporaso, J. G., Lauber, C. L., Costello, E. K., Berg-Lyons, D., Gonzalez, A., Stombaugh, J., Knights, D., Gajer, P., Ravel, J., Fierer, N., Gordon, J. I., & Knight, R. (2011). Moving pictures of the human microbiome. Genome Biology, 12(5). https://doi.org/10.1186/gb-2011-12-5-r50

12. Capuzzo, E., Lynam, C. P., Barry, J., Stephens, D., Forster, R. M., Greenwood, N., McQuatters-Gollop, A., Silva, T., van Leeuwen, S. M., & Engelhard, G. H. (2018). A decline in primary production in the North Sea over 25 years, associated with reductions in zooplankton abundance and fish stock recruitment. Global Change Biology, 24(1), e352–e364. https://doi.org/10.1111/gcb.13916

13. Carradec, Q., Poulain, J., Boissin, E., Hume, B. C., Voolstra, C. R., Ziegler, M., Engelen, S., Cruaud, C., Planes, S., & Wincker, P. (2020). A framework for in situ molecular characterization of coral holobionts using nanopore sequencing. BioRxiv Genomics, 10(1), 1–25. https://doi.org/10.1101/2020.05.25.071951

14. Chamberlain, S. A., & Szöcs, E. (2013). Taxize: Taxonomic search and retrieval in R. F1000Research, 2. https://doi.org/10.12688/f1000research.2-191.v2

15. Claudet, J. (2018). Six conditions under which MPAs might not appear effective (when they are). ICES Journal of Marine Science, 75(3), 1172–1174. https://doi.org/10.1093/icesjms/fsx074

16. Cole, J. R., Chai, B., Marsh, T. L., Farris, R. J., Wang, Q., Kulam, S. A., Chandra, S., McGarrell, D. M., Schmidt, T. M., Garrity, G. M., & Tiedje, J. M. (2003). The Ribosomal Database Project (RDP-II): previewing a new autoaligner that allows regular updates and the new prokaryotic taxonomy. Nucleic Acids Research, 31(1), 442–443. https://doi.org/10.1093/NAR/GKG039

17. Cordier, T., Frontalini, F., Cermakova, K., Apothéloz-Perret-Gentil, L., Treglia, M., Scantamburlo, E., Bonamin, V., & Pawlowski, J. (2019). Multi-marker eDNA metabarcoding survey to assess the environmental impact of three offshore gas platforms in the North Adriatic Sea (Italy). Marine Environmental Research, 146, 24–34. https://doi.org/10.1016/J.MARENVRES.2018.12.009

18. Couce, E., Schratzberger, M., & Engelhard, G. H. (2020). Reconstructing three decades of total international trawling effort in the North Sea. Earth System Science Data, 12(1), 373–386. https://doi.org/10.5194/essd-12-373-2020

19. Daan, N., Gislason, H., Pope, J. G., & Rice, J. C. (2005). Changes in the North Sea fish community: Evidence of indirect effects of fishing? ICES Journal of Marine Science, 62(2), 177–188. https://doi.org/10.1016/j.icesjms.2004.08.020

20. De Coster, W., D’Hert, S., Schultz, D. T., Cruts, M., & Van Broeckhoven, C. (2018). NanoPack: Visualizing and processing long-read sequencing data. Bioinformatics, 34(15), 2666–2669. https://doi.org/10.1093/bioinformatics/bty149

21. Deagle, B. E., Kirkwood, R., & Jarman, S. N. (2009). Analysis of Australian fur seal diet by pyrosequencing prey DNA in faeces. Molecular Ecology, 18(9), 2022–2038. https://doi.org/10.1111/j.1365-294X.2009.04158.x

22. Degraer, S., Carey, D. A., Coolen, J. W. P., Hutchison, Z. L., Kerckhof, F., Rumes, B., & Vanaverbeke, J. (2020). OFFSHORE WIND FARM ARTIFICIAL REEFS AFFECT ECOSYSTEM STRUCTURE AND FUNCTIONING. SPECIAL ISSUE ON UNDERSTANDING THE EFFECTS OF OFFSHORE WIND ENERGY DEVELOPMENT ON FISHERIES, 33(4), 48–57. https://doi.org/10.2307/26965749

23. Deiner, K., Bik, H. M., Mächler, E., Seymour, M., Lacoursière-Roussel, A., Altermatt, F., Creer, S., Bista, I., Lodge, D. M., de Vere, N., Pfrender, M. E., & Bernatchez, L. (2017). Environmental DNA metabarcoding: Transforming how we survey animal and plant communities. In Molecular Ecology (Vol. 26, Issue 21, pp. 5872–5895). Blackwell Publishing Ltd. https://doi.org/10.1111/mec.14350

24. Devloo-Delva, F., Huerlimann, R., Chua, G., Matley, J. K., Heupel, M. R., Simpfendorfer, C. A., & Maes, G. E. (2019). How does marker choice affect your diet analysis: comparing genetic markers and digestion levels for diet metabarcoding of tropical-reef piscivores. Marine and Freshwater Research, 70(1), 8. https://doi.org/10.1071/MF17209

25. DiBattista, J. D., Coker, D. J., Sinclair-Taylor, T. H., Stat, M., Berumen, M. L., & Bunce, M. (2017). Assessing the utility of eDNA as a tool to survey reef-fish communities in the Red Sea. In Coral Reefs (Vol. 36, Issue 4, pp. 1245–1252). Springer Verlag. https://doi.org/10.1007/s00338-017-1618-1

26. Didderen, K., Lengkeek, W., Bergsma, J. H., & Dongen, U. Van. (2019). WWF & ARK Borkum Reef Ground oyster pilot. September.

27. Doi, H., Fukaya, K., Oka, S. ichiro, Sato, K., Kondoh, M., & Miya, M. (2019). Evaluation of detection probabilities at the water-filtering and initial PCR steps in environmental DNA metabarcoding using a multispecies site occupancy model. Scientific Reports, 9(1), 1–8. https://doi.org/10.1038/s41598-019-40233-1

28. Edgar, R. C. (2004). MUSCLE: A multiple sequence alignment method with reduced time and space complexity. BMC Bioinformatics, 5. https://doi.org/10.1186/1471-2105-5-113

29. Egeter, B., Veríssimo, J., Lopes-Lima, M., Chaves, C., Pinto, J., Riccardi, N., Beja, P., & Fonseca, N. A. (2020). Speeding up the detection of invasive aquatic species using environmental DNA and nanopore sequencing. BioRxiv, 326(1), 2020.06.09.142521. https://doi.org/10.1101/2020.06.09.142521

30. Eggleton, J. D., Depestele, J., Kenny, A. J., Bolam, S. G., & Garcia, C. (2018). How benthic habitats and bottom trawling affect trait composition in the diet of seven demersal and benthivorous fish species in the North Sea. Journal of Sea Research, 142, 132–146. https://doi.org/10.1016/j.seares.2018.09.013

31. Evans, N. T., Li, Y., Renshaw, M. A., Olds, B. P., Deiner, K., Turner, C. R., Jerde, C. L., Lodge, D. M., Lamberti, G. A., & Pfrender, M. E. (2017). Fish community assessment with eDNA metabarcoding: Effects of sampling design and bioinformatic filtering. Canadian Journal of Fisheries and Aquatic Sciences, 74(9), 1362–1374. https://doi.org/10.1139/cjfas-2016-0306

32. Evans, N. T., Olds, B. P., Renshaw, M. A., Turner, C. R., Li, Y., Jerde, C. L., Mahon, A. R., Pfrender, M. E., Lamberti, G. A., & Lodge, D. M. (2016). Quantification of mesocosm fish and amphibian species diversity via environmental DNA metabarcoding. Molecular Ecology Resources, 16(1), 29–41. https://doi.org/10.1111/1755-0998.12433

33. Federhen, S. (2012). The NCBI Taxonomy database. Nucleic Acids Research, 40(D1), D136–D143. https://doi.org/10.1093/nar/gkr1178

34. Ficetola, G. F., Miaud, C., Pompanon, F., & Taberlet, P. (2008). Species detection using environmental DNA from water samples. Biology Letters, 4(4), 423–425. https://doi.org/10.1098/rsbl.2008.0118

35. Goodwin, K. D., Thompson, L. R., Duarte, B., Kahlke, T., Thompson, A. R., Marques, J. C., & Caçador, I. (2017). DNA sequencing as a tool to monitor marine ecological status. In Frontiers in Marine Science (Vol. 4, Issue MAY). Frontiers Media S. A. https://doi.org/10.3389/fmars.2017.00107

36. Heikema, A. P., Horst-Kreft, D., Boers, S. A., Jansen, R., Hiltemann, S. D., De Koning, W., Kraaij, R., De Ridder, M. A. J., Van Houten, C. B., Bont, L. J., Stubbs, A. P., & Hays, J. P. (2020). Comparison of illumina versus nanopore 16S rRNA gene sequencing of the human nasal microbiota. Mdpi.Com, 11. https://doi.org/10.3390/genes11091105

37. Hestetun, J. T., Bye-Ingebrigtsen, E., Nilsson, R. H., Glover, A. G., Johansen, P. O., & Dahlgren, T. G. (2020). Significant taxon sampling gaps in DNA databases limit the operational use of marine macrofauna metabarcoding. Marine Biodiversity, 50(5), 1–9. https://doi.org/10.1007/s12526-020-01093-5

38. Hofstede, R. Ter, Hiddink, J. G., & Rijnsdorp, A. D. (2010). Regional warming changes fish species richness in the eastern North Atlantic Ocean. Marine Ecology Progress Series, 414(August), 1–9. https://doi.org/10.3354/meps08753

39. Huang, Y., Niu, B., Gao, Y., Fu, L., & Li, W. (2010). CD-HIT Suite: A web server for clustering and comparing biological sequences. Bioinformatics, 26(5), 680–682. https://doi.org/10.1093/bioinformatics/btq003

40. Hunter, M. E., Ferrante, J. A., Meigs-Friend, G., & Ulmer, A. (2019). Improving eDNA yield and inhibitor reduction through increased water volumes and multi-filter isolation techniques. Scientific Reports, 9(1), 1–9. https://doi.org/10.1038/s41598-019-40977-w

41. Jain, M., Fiddes, I. T., Miga, K. H., Olsen, H. E., Paten, B., & Akeson, M. (2015). Improved data analysis for the MinION nanopore sequencer. Nature Methods, 12(4), 351–356. https://doi.org/10.1038/nmeth.3290

42. Jin, J., Chang, M., Cheong, Y., Ip, A., Bauman, A. G., & Huang, D. (2020). MinION-in-ARMS: Nanopore Sequencing To Expedite Barcoding Of Specimen-Rich Macrofaunal Samples From Autonomous Reef 2 Monitoring Structures 3 4. BioRxiv, 2020.03.30.009654. https://doi.org/10.1101/2020.03.30.009654

43. Johnson, J. S., Spakowicz, D. J., Hong, B. Y., Petersen, L. M., Demkowicz, P., Chen, L., Leopold, S. R., Hanson, B. M., Agresta, H. O., Gerstein, M., Sodergren, E., & Weinstock, G. M. (2019). Evaluation of 16S rRNA gene sequencing for species and strain-level microbiome analysis. Nature Communications, 10(1), 1–11. https://doi.org/10.1038/s41467-019-13036-1

44. Kamermans P., B. Walles, M. Kraan, L.A. van Duren, F. Kleissen & T.M. van der Have, A.C. Smaal, M. Poelman (2018) Offshore wind farms as potential locations for flat oyster (Ostrea edulis) restoration in the Dutch North Sea. Sustainability 10, 3942; doi:10.3390/su10113942Katoh, K., & Standley, D. M. (2013). MAFFT Multiple Sequence Alignment Software Version 7: Improvements in Performance and Usability. Molecular Biology and Evolution, 30(4), 772–780. https://doi.org/10.1093/molbev/mst010

45. Kearse, M., Moir, R., Wilson, A., Stones-Havas, S., Cheung, M., Sturrock, S., Buxton, S., Cooper, A., Markowitz, S., Duran, C., Thierer, T., Ashton, B., Meintjes, P., & Drummond, A. (2012). Geneious Basic: An integrated and extendable desktop software platform for the organization and analysis of sequence data. Bioinformatics, 28(12), 1647–1649. https://doi.org/10.1093/bioinformatics/bts199

46. Kelly, R. P., Shelton, A. O., & Gallego, R. (2019). Understanding PCR Processes to Draw Meaningful Conclusions from Environmental DNA Studies. Scientific Reports, 9(1). https://doi.org/10.1038/s41598-019-48546-x

47. Kim, D., Song, L., Breitwieser, F. P., & Salzberg, S. L. (2016). Centrifuge: Rapid and sensitive classification of metagenomic sequences. Genome Research, 26(12), 1721–1729. https://doi.org/10.1101/gr.210641.116

48. Krehenwinkel, H., Pomerantz, A., & Prost, S. (2019). Genetic biomonitoring and biodiversity assessment using portable sequencing technologies: Current uses and future directions. Genes, 10(11). https://doi.org/10.3390/genes10110858

49. Lacoursière-Roussel, A., Côté, G., Leclerc, V., & Bernatchez, L. (2016). Quantifying relative fish abundance with eDNA: a promising tool for fisheries management. Journal of Applied Ecology, 53(4), 1148–1157. https://doi.org/10.1111/1365-2664.12598

50. Leray, M., Yang, J. Y., Meyer, C. P., Mills, S. C., Agudelo, N., Ranwez, V., Boehm, J. T., & Machida, R. J. (2013). A new versatile primer set targeting a short fragment of the mitochondrial COI region for metabarcoding metazoan diversity: Application for characterizing coral reef fish gut contents. Frontiers in Zoology, 10(1). https://doi.org/10.1186/1742-9994-10-34

51. Li, H. (2018). Minimap2: Pairwise alignment for nucleotide sequences. Bioinformatics, 34(18), 3094– 3100. https://doi.org/10.1093/bioinformatics/bty191

52. Li, W., Jaroszewski, L., & Godzik, A. (2002). Tolerating some redundancy significantly speeds up clustering of large protein databases. Bioinformatics, 18(1), 77–82. https://doi.org/10.1093/bioinformatics/18.1.77

53. Manel, S., Guerin, P. E., Mouillot, D., Blanchet, S., Velez, L., Albouy, C., & Pellissier, L. (2020). Global determinants of freshwater and marine fish genetic diversity. Nature Communications, 11(1), 1–9. https://doi.org/10.1038/s41467-020-14409-7

54. Martin, M. (2011). Cutadapt removes adapter sequences from high-throughput sequencing reads. EMBnet.Journal, 17(1), 10. https://doi.org/10.14806/EJ.17.1.200

55. Mateos-Rivera, A., Skern-Mauritzen, R., Dahle, G., Sundby, S., Mozfar, B., Thorsen, A., Wehde, H., & Krafft, B. A. (2020). Comparison of visual and molecular taxonomic methods to identify ichthyoplankton in the North Sea. Limnology and Oceanography: Methods. https://doi.org/10.1002/lom3.10387

56. Matsuo, Y., Komiya, S., Yasumizu, Y., Yasuoka, Y., Mizushima, K., Takagi, T., Kryukov, K., Fukuda, A., Morimoto, Y., Naito, Y., Okada, H., Bono, H., Nakagawa, S., & Hirota, K. (2021). Full-length 16S rRNA gene amplicon analysis of human gut microbiota using MinION^TM^ nanopore sequencing confers species-level resolution. BMC Microbiology, 21(1). https://doi.org/10.1186/s12866-021-02094-5

57. McGee, K. M., Robinson, C. V., & Hajibabaei, M. (2019). Gaps in DNA-Based Biomonitoring Across the Globe. Frontiers in Ecology and Evolution, 7, 337. https://doi.org/10.3389/fevo.2019.00337

58. McMurdie, P. J., & Holmes, S. (2013). phyloseq: An R Package for Reproducible Interactive Analysis and Graphics of Microbiome Census Data. PLoS ONE, 8(4), e61217. https://doi.org/10.1371/journal.pone.0061217

59. Miya, M., Sato, Y., Fukunaga, T., Sado, T., Poulsen, J. Y., Sato, K., Minamoto, T., Yamamoto, S., Yamanaka, H., Araki, H., Kondoh, M., & Iwasaki, W. (2015a). MiFish, a set of universal PCR primers for metabarcoding environmental DNA from fishes: Detection of more than 230 subtropical marine species. Royal Society Open Science, 2(7), 150088. https://doi.org/10.1098/rsos.150088

60. Miya, M., Sato, Y., Fukunaga, T., Sado, T., Poulsen, J. Y., Sato, K., Minamoto, T., Yamamoto, S., Yamanaka, H., Araki, H., Kondoh, M., & Iwasaki, W. (2015b). MiFish, a set of universal PCR primers for metabarcoding environmental DNA from fishes: Detection of more than 230 subtropical marine species. Royal Society Open Science, 2(7). https://doi.org/10.1098/rsos.150088

61. Morey, K. C., Bartley, T. J., & Hanner, R. H. (2020). Validating environmental DNA metabarcoding for marine fishes in diverse ecosystems using a public aquarium. Environmental DNA, 2(3), 330–342. https://doi.org/10.1002/edn3.76

62. O’Brien, A. L., Dafforn, K. A., Chariton, A. A., Johnston, E. L., & Mayer-Pinto, M. (2019). After decades of stressor research in urban estuarine ecosystems the focus is still on single stressors: A systematic literature review and meta-analysis. In Science of the Total Environment (Vol. 684, pp. 753–764). Elsevier B.V. https://doi.org/10.1016/j.scitotenv.2019.02.131

63. Ratnasingham, S., & Hebert, P. D. N. (2007). BOLD: The Barcode of Life Data System: Barcoding. Molecular Ecology Notes, 7(3), 355–364. https://doi.org/10.1111/j.1471-8286.2007.01678.x

64. Reiss, H., Degraer, S., Duineveld, G. C. A., Kröncke, I., Aldridge, J., Craeymeersch, J. A., Eggleton, J. D., Hillewaert, H., Lavaleye, M. S. S., Moll, A., Pohlmann, T., Rachor, E., Robertson, M., Vanden Berghe, E., Van Hoey, G., & Rees, H. L. (2010). Spatial patterns of infauna, epifauna, and demersal fish communities in the North Sea. ICES Journal of Marine Science, 67(2), 278–293. https://doi.org/10.1093/icesjms/fsp253

65. Riaz, T., Shehzad, W., Viari, A., Pompanon, F., Taberlet, P., & Coissac, E. (2011). EcoPrimers: Inference of new DNA barcode markers from whole genome sequence analysis. Nucleic Acids Research, 39(21), e145. https://doi.org/10.1093/nar/gkr732

66. Rodríguez-Pérez, H., Ciuffreda, L., & Flores, C. (2020). NanoCLUST: a species-level analysis of 16S rRNA nanopore sequencing data. Bioinformatics. https://doi.org/10.1093/bioinformatics/btaa900

67. Ruppert, K. M., Kline, R. J., & Rahman, M. S. (2019a). Past, present, and future perspectives of environmental DNA (eDNA) metabarcoding: A systematic review in methods, monitoring, and applications of global eDNA. In Global Ecology and Conservation (Vol. 17, p. e00547). Elsevier B.V. https://doi.org/10.1016/j.gecco.2019.e00547

68. Ruppert, K. M., Kline, R. J., & Rahman, M. S. (2019b). Past, present, and future perspectives of environmental DNA (eDNA) metabarcoding: A systematic review in methods, monitoring, and applications of global eDNA. In Global Ecology and Conservation (Vol. 17, p. e00547). Elsevier B.V. https://doi.org/10.1016/j.gecco.2019.e00547

69. Santos, A., van Aerle, R., Barrientos, L., & Martinez-Urtaza, J. (2020a). Computational methods for 16S metabarcoding studies using Nanopore sequencing data. In Computational and Structural Biotechnology Journal (Vol. 18, pp. 296–305). Elsevier B.V. https://doi.org/10.1016/j.csbj.2020.01.005

70. Santos, A., van Aerle, R., Barrientos, L., & Martinez-Urtaza, J. (2020b). Computational methods for 16S metabarcoding studies using Nanopore sequencing data. In Computational and Structural Biotechnology Journal (Vol. 18, pp. 296–305). Elsevier B.V. https://doi.org/10.1016/j.csbj.2020.01.005

71. Sard, N. M., Herbst, S. J., Nathan, L., Uhrig, G., Kanefsky, J., Robinson, J. D., & Scribner, K. T. (2019). Comparison of fish detections, community diversity, and relative abundance using environmental DNA metabarcoding and traditional gears. Environmental DNA, 1(4), 368–384. https://doi.org/10.1002/edn3.38

72. Sassoubre, L. M., Yamahara, K. M., Gardner, L. D., Block, B. A., & Boehm, A. B. (2016). Quantification of Environmental DNA (eDNA) Shedding and Decay Rates for Three Marine Fish. Environmental Science and Technology, 50(19), 10456–10464. https://doi.org/10.1021/acs.est.6b03114

73. Schon, E. A. (2000). Mitochondrial genetics and disease. In Trends in Biochemical Sciences (Vol. 25, Issue 11, pp. 555–560). Trends Biochem Sci. https://doi.org/10.1016/S0968-0004(00)01688-1

74. Schroeter, J. C., Maloy, A. P., Rees, C. B., & Bartron, M. L. (2020). Fish mitochondrial genome sequencing: expanding genetic resources to support species detection and biodiversity monitoring using environmental DNA. Conservation Genetics Resources, 12(3), 433–446. https://doi.org/10.1007/s12686-019-01111-0

75. Seymour, M., Durance, I., Cosby, B. J., Ransom-Jones, E., Deiner, K., Ormerod, S. J., Colbourne, J. K., Wilgar, G., Carvalho, G. R., de Bruyn, M., Edwards, F., Emmett, B. A., Bik, H. M., & Creer, S. (2018). Acidity promotes degradation of multi-species environmental DNA in lotic mesocosms. Communications Biology, 1(1). https://doi.org/10.1038/s42003-017-0005-3

76. Shin, J., Lee, S., Go, M. J., Lee, S. Y., Kim, S. C., Lee, C. H., & Cho, B. K. (2016). Analysis of the mouse gut microbiome using full-length 16S rRNA amplicon sequencing. Scientific Reports, 6. https://doi.org/10.1038/srep29681

77. Shu, L., Ludwig, A., & Peng, Z. (2020). Standards for methods utilizing environmental dna for detection of fish species. In Genes (Vol. 11, Issue 3, p. 296). MDPI AG. https://doi.org/10.3390/genes11030296

78. Singer, G. A. C., Fahner, N. A., Barnes, J. G., McCarthy, A., & Hajibabaei, M. (2019). Comprehensive biodiversity analysis via ultra-deep patterned flow cell technology: a case study of eDNA metabarcoding seawater. Scientific Reports, 9(1), 1–12. https://doi.org/10.1038/s41598-019-42455-9

79. Srivathsan, A., Baloğlu, B., Wang, W., Tan, W. X., Bertrand, D., Ng, A. H. Q., Boey, E. J. H., Koh, J. J. Y., Nagarajan, N., & Meier, R. (2018). A MinION^TM^-based pipeline for fast and cost-effective DNA barcoding. Molecular Ecology Resources, 18(5), 1035–1049. https://doi.org/10.1111/1755-0998.12890

80. Srivathsan, A., Lee, L., Katoh, K., Hartop, E., Kutty, S. N., Wong, J., Yeo, D., & Meier, R. (2021). ONTbarcoder and MinION barcodes aid biodiversity discovery and identification by everyone, for everyone. BMC Biology, 19(1). https://doi.org/10.1186/S12915-021-01141-X

81. Taberlet, P., Bonin, A., Zinger, L., & Coissac, E. (2018). Environmental DNA: For biodiversity research and monitoring. In Environmental DNA: For Biodiversity Research and Monitoring. https://doi.org/10.1093/oso/9780198767220.001.0001

82. Taberlet, P., Coissac, E., Pompanon, F., Brochmann, C., & Willerslev, E. (2012). Towards next-generation biodiversity assessment using DNA metabarcoding. Molecular Ecology, 21(8), 2045–2050. https://doi.org/10.1111/j.1365-294X.2012.05470.x

83. Tan, G., Opitz, L., Schlapbach, R., & Rehrauer, H. (2019). Long fragments achieve lower base quality in Illumina paired-end sequencing. Scientific Reports, 9(1), 1–7. https://doi.org/10.1038/s41598-019-39076-7

84. Teletchea, F. (2009). Molecular identification methods of fish species: Reassessment and possible applications. Reviews in Fish Biology and Fisheries, 19(3), 265–293. https://doi.org/10.1007/s11160-009-9107-4

85. Thalinger, B., Wolf, E., Traugott, M., & Wanzenböck, J. (2019). Monitoring spawning migrations of potamodromous fish species via eDNA. Scientific Reports, 9(1), 1–11. https://doi.org/10.1038/s41598-019-51398-0

86. Thomsen, P. F., Kielgast, J., Iversen, L. L., Møller, P. R., Rasmussen, M., & Willerslev, E. (2012). Detection of a Diverse Marine Fish Fauna Using Environmental DNA from Seawater Samples. PLoS ONE, 7(8). https://doi.org/10.1371/journal.pone.0041732

87. Thomsen, P. F., & Willerslev, E. (2015). Environmental DNA – An emerging tool in conservation for monitoring past and present biodiversity. Biological Conservation, 183, 4–18. https://doi.org/10.1016/J.BIOCON.2014.11.019

88. Truelove, N. K., Andruszkiewicz, E. A., & Block, B. A. (2019). A rapid environmental DNA method for detecting white sharks in the open ocean. Methods in Ecology and Evolution, 10(8), 1128–1135. https://doi.org/10.1111/2041-210X.13201

89. van der Loos, L. M., & Nijland, R. (2020). Biases in bulk: DNA metabarcoding of marine communities and the methodology involved. Molecular Ecology. https://doi.org/10.1111/mec.15592

90. Van Der Reis, A. L., Lynnath, |, Beckley, E., Olivar, | M Pilar, & Jeffs, A. G. (2023). Nanopore short-read sequencing: A quick, cost-effective and accurate method for DNA metabarcoding. Wiley Online Library. https://doi.org/10.1002/edn3.374

91. Vaser, R., Sović, I., Nagarajan, N., & Šikić, M. (2017). Fast and accurate de novo genome assembly from long uncorrected reads. Genome Research, 27(5), 737–746. https://doi.org/10.1101/gr.214270.116

92. Vaser, R., Sović, I., Sović, S., Nagarajan, N., Šikić1, M., & Šikić1, Š. (2017). Fast and accurate de novo genome assembly from long uncorrected reads. Genome.Cshlp.Org. https://doi.org/10.1101/gr.214270.116

93. Voorhuijzen-Harink, M. M., Hagelaar, R., van Dijk, J. P., Prins, T. W., Kok, E. J., & Staats, M. (2019). Toward on-site food authentication using nanopore sequencing. Food Chemistry: X, 2, 100035. https://doi.org/10.1016/j.fochx.2019.100035

94. Wang, S., Yan, Z., Hänfling, B., Zheng, X., Wang, P., Fan, J., & Li, J. (2021). Methodology of fish eDNA and its applications in ecology and environment. In Science of the Total Environment (Vol. 755, p. 142622). Elsevier B.V. https://doi.org/10.1016/j.scitotenv.2020.142622

95. Zhang, S., Zhao, J., & Yao, M. (2020). A comprehensive and comparative evaluation of primers for metabarcoding eDNA from fish. Methods in Ecology and Evolution. https://doi.org/10.1111/2041-210x.13485

